# Reconstructing the human enhancer RNA transcriptome

**DOI:** 10.64898/2026.03.02.709149

**Authors:** Natalia Benova, Rene Kuklinkova, Emanuela Ibenye, James R. Boyne, Chinedu A. Anene

**Affiliations:** Centre for Biomedical Science Research, School of Health, Leeds Beckett University, Leeds, LS1 3HE, United Kingdom

**Keywords:** Enhancer RNAs, Transcript-resolved eRNA models, RNA splicing

## Abstract

Transcript-resolved models of RNA enable functional interrogation of RNA biology by linking processing, structure, localisation, and regulatory interactions to specific RNA molecules. Across coding and noncoding transcriptomes, such models have been essential for defining RNA-level mechanisms relevant to physiology and disease. Enhancer RNAs (eRNAs), however, remain largely characterised without transcript-level definitions, and no widely adopted transcript-resolved reference exists, limiting investigation of how individual eRNAs are processed, localised, and participate in transcriptional regulation or their emerging post-transcriptional functions. Here, we reconstruct a transcript-resolved catalogue of human eRNAs by pan-transcriptome assembly across diverse tissues, cell types and compartments, defining 36,536 transcripts, including a subset with multi-exonic structure.

We show that eRNA splice junctions are reproducible features that exhibit cell-type specificity, subcellular localisation bias, and sensitivity to spliceosome perturbation. In perturbation experiments, eRNA splice junction usage responded to SF3B1 mutation, nuclear–cytoplasmic partitioning, and pharmacological inhibition of RNA export, demonstrating regulation across multiple layers of RNA biology. In head and neck squamous cell carcinoma, a subset of these junctions showed altered usage between tumour and matched normal tissue, indicating that processing varies in disease contexts. Across three validation contexts, nearly one-fifth of reconstructed junctions were detectable, with some showing regulated usage, supporting biological reproducibility.

Motivated by these observations, we provide both the GTF annotation and junctions BED file, as a framework for studying eRNAs, enabling RNA-centric investigation of their potential functions. The annotations have been incorporated into the eRNAkit database, available at https://github.com/AneneLab/eRNAkit.

## Introduction

Active enhancers are marked by transcription that produces enhancer RNAs (eRNAs) ^1^, yet the functions of these RNA molecules remain poorly defined. This partly reflects how eRNAs are defined: they are mostly annotated as genomic coordinates or nascent transcription signals rather than transcript structures ^2–4^. Consequently, eRNAs are interpreted primarily as indicators of enhancer activation state rather than as a set of RNA species with defined boundaries and functions.

Nevertheless, growing evidence suggests that eRNAs comprise a heterogeneous population of RNA molecules, implying that individual eRNA species may carry functions at the RNA level beyond their role in transcriptional regulation ^3,5^. Many eRNAs are retained in the nucleus, where functional studies link them to chromatin organisation, enhancer–promoter communication, and transcriptional control ^6^. At the same time, multiple datasets detect subsets in cytoplasmic and organelle-associated fractions, and several studies report trans-acting and post-transcriptional effects that are difficult to reconcile with strictly local enhancer–promoter models ^5,7,8^. These observations suggest that at least a subset of eRNAs act as functional RNA molecules rather than solely reflecting enhancer activity. However, without transcript-level definitions, it remains unclear which specific RNA species mediate these effects. Defining enhancer-derived transcripts is therefore essential to investigate how individual eRNAs are processed, localised, structurally organised, and engaged by RNA-binding proteins.

This conceptual limitation directly affects how eRNA function is investigated. Many current eRNAs studies that use RNA interference select targets based on enhancer genomic coordinates, implicitly treating eRNAs as single transcriptional units rather than as structured RNA molecules. Yet evidence that some eRNAs undergo processing, stabilisation, and export beyond their sites of synthesis indicates that distinct transcripts arising from the same enhancer may possess divergent functions, analogous to other processed long noncoding RNAs ^1^. If eRNAs operate at the level of RNA molecules rather than genomic loci, coordinate-based targeting risks missing RNA-specific regulatory effects and may confound interpretation when multiple transcripts from the same enhancer are functionally non-equivalent.

Here, we assemble a pan-transcriptome of eRNAs across diverse human tissues and cell types, defining them at the level of RNA molecules rather than genomic coordinates. This reconstruction resolves enhancer transcription into structured eRNA transcripts with defined boundaries and isoforms and shows that some loci can generate strand-resolved multi-exonic RNAs with features consistent with RNA processing. By providing transcript-level models and cross-indexing them to existing enhancer annotations, this resource establishes a transcript-centric framework for studying eRNA function and enables isoform-aware analyses and targeting strategies across the community.

## Results

### Transcript-level reconstruction of human enhancer RNAs

We reconstructed enhancer-derived transcripts across diverse human tissues and cell lines using a pan-transcriptome assembly of stranded, high depth Encode RNA-Seq datasets spanning (n=121, Table S1). To focus reconstruction on enhancer transcription while preserving transcript resolution, we used our previously curated eRNA coordinates as a reference framework (n = 48,766) ^7^. RNA-Seq reads were filtered to retain only those overlapping these regions, enriching for eRNAs while minimising signal from gene bodies and promoters (Methods).

Because robust reconstruction of low-abundance eRNAs requires aggregation of transcriptional evidence, we first evaluated two pooling strategies in a pilot analysis on a small subset of samples (n = 5), comparing (i) transcript assembly from pooled coverage and (ii) independent per-sample assembly followed by model. Evaluation against annotated genes and transcripts showed that assembly from merged coverage recover more known transcript structures than per sample assembly, consistent with prior pooled-signal approaches used to define enhancer transcription landscapes ^1^.

Next, we applied it to the filtered eRNA overlapping reads to reconstruct transcript models, with a minimum transcript length of 100 nt. Transcript lengths ranged from 100 nt to 99,000 nt (median 957 nt), with 95% shorter than 20,000 nt, consistent with previous reports that eRNAs are typically short (Fig. 1a). We therefore applied conservative filters to remove models unlikely to reflect continuous transcription, excluding transcripts longer than 100,000 nt and eliminating 463 models together with their associated exons (1.8%). We next reassessed overlap with annotated genes, since reconstruction across pooled samples can extend transcript boundaries beyond the original enhancer coordinates. Comparison with RefSeq annotations identified 8,620 gene-intersecting transcripts, which we excluded to minimise contamination from gene-associated transcription. Compared to the retained models, these excluded overlaps were enriched at both extremes of the length distribution (Fig. 1b).

**Figure 1.**
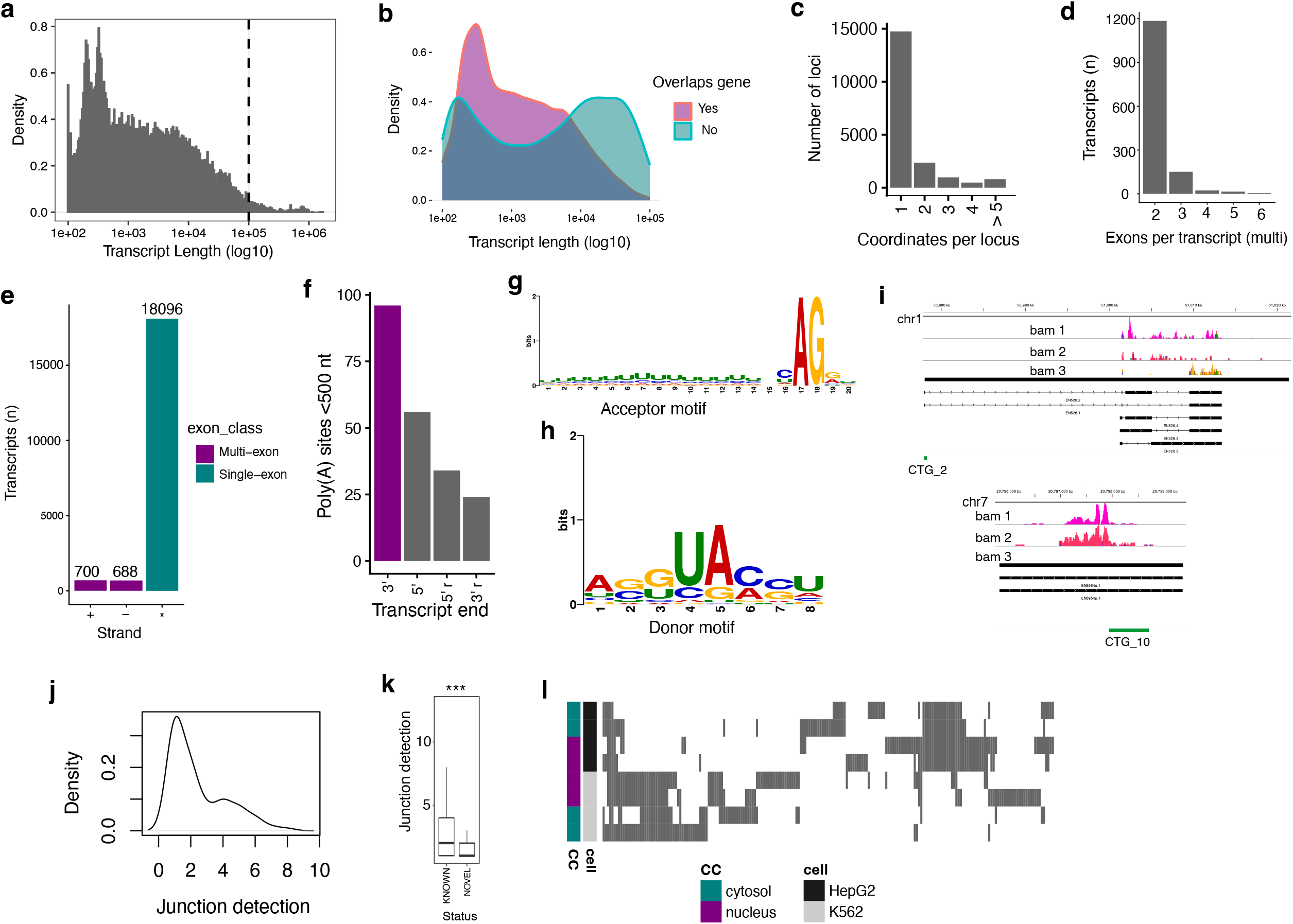
Transcript-level reconstruction and structural features of human enhancer RNAs. (a) Length (nt) distribution of reconstructed eRNA transcripts across the assembled catalogue. Vertical black line indicates excluded long transcripts. (b) Length (nt) distribution of transcripts removed due to overlap with annotated gene bodies compared with retained eRNA transcripts. (c) Bar plot showing how reference curated enhancer coordinates merged into transcript models after assembly. (d) Distribution of exon number per transcript across reconstructed eRNAs. Only multi exonic transcript are shown. (e) Strand distribution of multi-exonic transcripts. (f) Distance of reconstructed transcript termini to annotated polyadenylation sites, shown for strand-consistent (3’ and 5’) and strand-reversed (3’ r and 5’ r) controls. (g–h) De novo motif discovery at reconstructed splice junctions for (g) splice acceptor sites and (h) splice donor sites derived from reconstructed junctions. (i) Overlap of previously reported eRNA ORFs with reconstructed transcript models, illustrating isoform-specific compatibility (top panel) and block version (bottom panel). (j) Number of samples supporting each reconstructed junction in the assembly subset. (k) Comparison of detection frequencies between reconstructed junctions (KNOWN) and additional junctions identified during counting (NOVEL). *** indicates p < 2.2e-16, Wilcoxon rank-sum test. (l) Heatmap showing presence or absence of reconstructed junctions across samples, with cell type and RNA fraction indicated.

The filtered models defined a human eRNA catalogue comprising 18,740 loci and 19,484 transcripts. This represents a substantial refinement over the starting set of 48,766 curated eRNA coordinates used as the reference framework. We found many adjacent reference coordinates that resolved into single multi-exonic (Fig. 1c), whereas some individual coordinates resolved into multiple reconstructed loci. Together, these patterns indicate that coordinate-based annotations can both fragment individual eRNA molecules and merge transcriptionally complex enhancer regions that produce more than one RNA species. To maintain compatibility with our original reference framework, each transcript coordinate was cross-indexed to our curated eRNAkit ID (GTF attribute: eRNAkit), enabling direct integration with previously defined expression, localisation, and activity data in eRNAKitDB.

### Human enhancer RNAs include structured, strand-resolved RNA molecules

Across the eRNA catalogue, most loci produced a single eRNA transcript (18,740 loci; 99.3%), whereas only a small fraction produced multiple isoforms (128 loci; 0.68%). At the exon level, most transcripts were unspliced (18,105 transcripts; 93%), while 1,379 contained multiple exons (7%). Among these, exon number remained low, with most multi-exonic transcripts comprising two or three exons (Fig. 1d) that were all were strand-resolved and distributed across both strands (603 + and 593 -, Fig. 1e). Consistent with this organisation, multi-exonic transcripts were substantially longer than unspliced transcripts (median 8.2 kb vs 0.8 kb; Wilcoxon rank-sum test, p < 2.2e-16), reflecting the broader length difference between strand-resolved and unstranded models (Kruskal–Wallis test, chi-squared = 1800.7, df = 2, p < 2.2e-16).

Together, these structural features indicate that strand-resolved eRNA molecules form defined RNA species, prompting us to test whether their reconstructed 3’ ends correspond to sites of RNA cleavage and termination. Because many eRNAs are not expected to be polyadenylated, we tested for enrichment, rather than absolute presence, of directionally consistent poly(A) sites at reconstructed transcript ends, using annotated cleavage sites from the PolyASite database ^9^, with strand-reversed relationships as orientation controls. 0.50% of 19,378 transcript 3′ termini were located within 500 nt of a directionally consistent polyadenylation site, compared with 0.29% of 19,357 matched 5′ ends from the same transcripts (Fig. 1f), representing a 1.7-fold enrichment at transcript termini (Fisher’s exact test, p = 1.45e-3). Strand-reversed controls were substantially more distant from poly(A) sites, with only 0.12% of opposite-strand 3′ ends felling within the same distance window, 4-fold fewer than true 3′ ends (Fisher’s exact test, p = 2.02e-11). 0.18% of opposite-strand 5′ ends fell within this window, again 2.8-fold fewer than true 3′ ends (Fisher’s exact test, p = 4.71e-8).

To assess whether reconstructed exon–intron junctions display canonical splice-site architecture, we analysed donor and acceptor sequences extracted from strand-resolved transcripts by MEME de novo motif discovery ^10^. Motifs were identified from 1,462 acceptor sequences and 1,443 donor sequences (Fig. g-h). Acceptor sites showed a highly significant enrichment of the canonical splice acceptor dinucleotide at the exon–intron boundary (Fig. 1g). The dominant motif was recovered in 1,458 junctions (E-value = 4.1e-1179 ; width = 20 nt), displaying a sharply defined AG signal with minimal positional dispersion. No additional significant motifs were detected, indicating strong conservation of acceptor structure across eRNA splice sites. Donor sites likewise displayed a significant consensus motif (Fig. 1h), detected in 1,441 junctions (E-value = 7.2e-875; width = 8 nt). In contrast to the acceptor signal, the donor motif extended across the exon–intron boundary rather than collapsing to a single conserved dinucleotide. This broader structure is consistent with experimentally defined human 5′ splice-site architecture, in which sequence preferences span both exon and intron and contribute to spliceosome recognition via U5 and U6 snRNA interactions ^11^.

The presence of strongly conserved acceptor signatures and structured donor consensus motifs demonstrates that reconstructed eRNA splice junctions exhibit canonical sequence features of functional splice sites. This prompted us to ask whether transcript-level reconstruction refines interpretation of eRNA-associated open reading frames (ORFs) reported in a recent study ^5^. Intersection of the published 53 ORF coordinates with our catalogue identified two overlaps (3.77%). One occurred at a locus producing multiple spliced isoforms, where only a subset of transcripts contained exon combinations compatible with the reported coding sequence (Fig. 1i), indicating that coding plausibility is isoform-dependent rather than locus-dependent. The second overlapped a single-exon transcript at a locus lacking confident strand resolution (Fig. 1i). These observations illustrate how transcript-resolved eRNA models provide structural context for interpreting coding potential by linking ORFs to specific RNA architectures rather than genomic regions alone.

### Reconstructed eRNA splice junctions are reproducible and sensitive to spliceosome perturbation

To assess whether reconstructed eRNA splice junctions represent biologically regulated transcript features, we first revisited a defined subset of the datasets underlying transcript assembly (n = 8), comprising poly(A)-selected nuclear and cytoplasmic RNA from K562 and HepG2. We quantified junction detection across these samples to determine whether eRNA junctions show consistent read support, cell-specific usage, or compartmental. Of the 1,538 reconstructed splice junctions, 206 (13.4%) were detected across the eight samples, with 56.3% observed in at least in two samples (116 of 206; Fig. 1j). Reconstructed transcript junctions were detected in significantly more samples than additional junctions identified during counting (Wilcoxon rank-sum test, p = 2.2e-16; Fig. 1k). Detection patterns segregated by both cell type and eRNA localisation: some junctions were restricted to either K562 or HepG2, others showed compartment bias within a cell type, and a smaller subset was detected broadly across samples (Fig. 1l). These patterns indicate that reconstructed eRNA splice junctions behave as structured, context-dependent transcript features rather than assembly artefacts, motivating validation in independent datasets with direct spliceosome perturbation.

We next examined splice-junction behaviour in an independent perturbation dataset from HeLa cells ^12^, which was not used for transcript assembly. This dataset includes treatment with two distinct spliceosome inhibitors: SpliceostatinA, which targets the SF3B complex and blocks spliceosome assembly, and Isoginkgetin, which inhibits spliceosome progression at an earlier stage of splice-site pairing. Across these samples, 69 reconstructed junctions showed detectable read support and were detected in significantly more samples than additional junctions identified during counting (Wilcoxon rank-sum test, P = 2.2e-16). Both Isoginkgetin and SpliceostatinA treatment were associated with an overall reduction in eRNA splice-junction signal relative to matched vehicle controls (Fig. 2a), consistent with impaired splicing. Sufficient coverage for differential usage testing was available for 10 junctions in the SpliceostatinA vs DMSO comparison and 14 in the Isoginkgetin vs Methanol pair.

**Figure 2.**
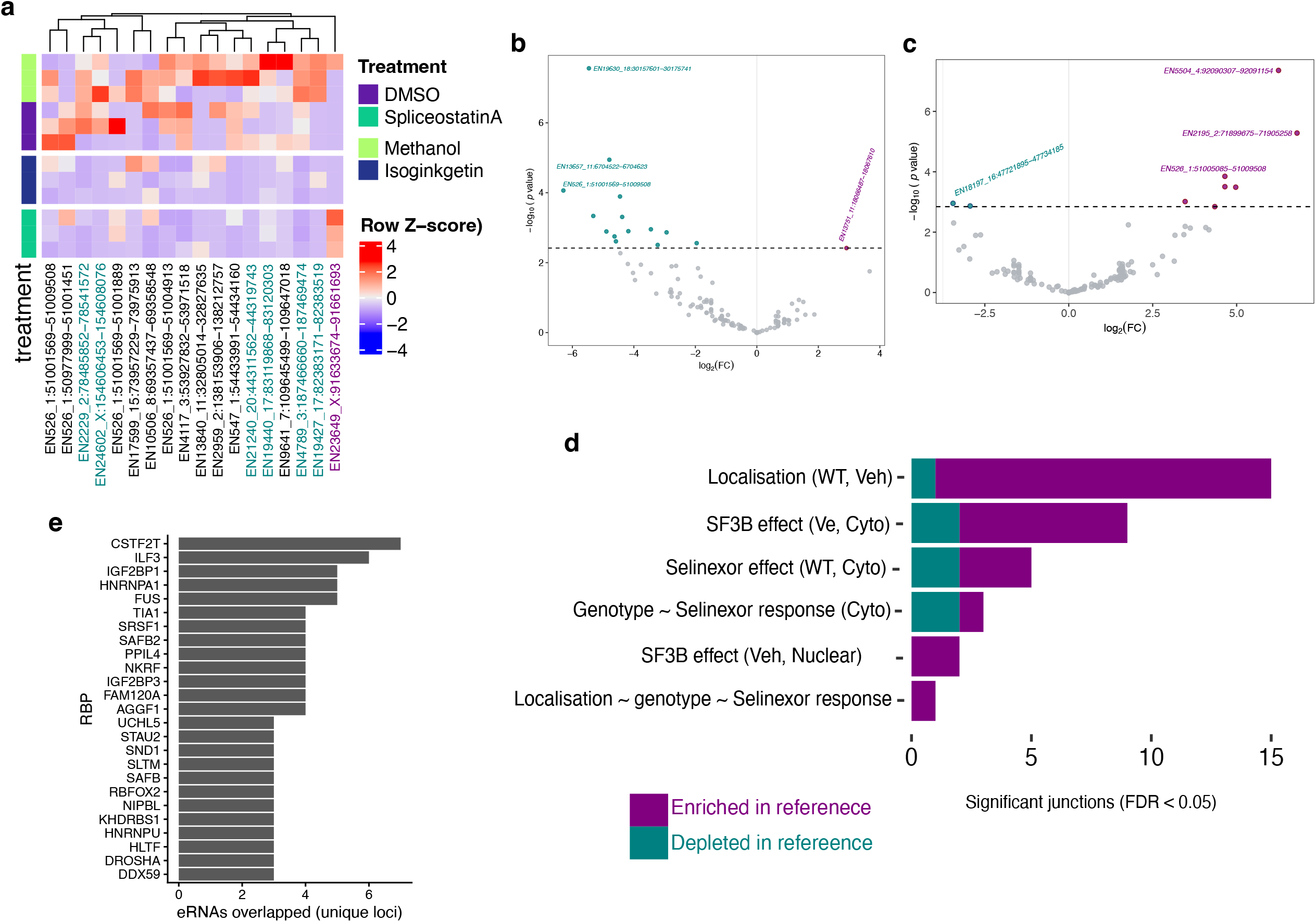
Detection and perturbation sensitivity of reconstructed enhancer RNA splice junctions. (a) Detection and differential usage of reconstructed splice junctions in HeLa cells treated with spliceosome inhibitors relative to vehicle controls (DMSO-SpliceostatinA and Methanol-Isoginkgetin). Within the plot, green coloured junctions are significant at FDR < 0.05 for Methanol-Isoginkgetin comparison and purple significant for DMSO-SpliceostatinA comparison. (b-c) Volcano plot of differential junction usage between nuclear and cytoplasmic RNA fractions in K562 cells, and between wild-type and SF3B1-mutant K562 cells under vehicle conditions. (d) Number of significant changes in junction usage across the full factorial analysis. Values in bracket indicate the background, WT – wildtype, Veh – vehicle control and Cyto – cytoplasm compartment. The ∼ indicates interactor term in the contrast. (e) RNA-binding proteins detected at reconstructed enhancer transcripts across ENCODE eCLIP datasets, summarised as number of loci with overlapping peaks.

SpliceostatinA altered usage of one junction, showing increased inclusion, whereas Isoginkgetin affected five junctions (5 of 14 tested; 35.7%), all showing reduced usage (Fig. 2a). Although these single-end libraries provide limited resolution for junction quantification, the consistent reduction in junction signal and the detectable differential responses demonstrate that reconstructed eRNA splice sites can respond to spliceosome perturbation in an independent cellular system.

We next analysed splice-junction behaviour in a larger independent perturbation dataset from K562 cells ^13^ to test whether reconstructed eRNA junctions respond to genetic and cellular perturbations. This dataset combines three orthogonal variables relevant to RNA processing: SF3B1 mutation (WT vs K666N), nuclear versus cytoplasmic RNA fractionation, and treatment with the nuclear-export inhibitor selinexor alongside matched DMSO controls. The design therefore allows simultaneous assessment of mutation effects, compartmental localisation, drug response, and their interactions on eRNA splice-junction usage in a controlled cellular background. Across the 24 paired-end samples, 456 of 1,538 reconstructed junctions were detected (29.7%), a substantially broader recovery than in the single-end HeLa dataset (69 of 1,538; 4.5%). As in the earlier analysis, reconstructed junctions were detected in significantly more samples than additional junctions identified during counting (Wilcoxon rank-sum test, P = 8e-06).

In this dataset, the strongest determinant of junction usage was eRNA localisation: in wild-type cells, 15 of 180 tested junctions (8.3%) differed between nuclear and cytoplasmic fractions (FDR < 0.05), with 14 showing higher usage in the cytoplasm and 1 showing higher usage in the nucleus, indicating that eRNA splice-junction usage varies with RNA localisation (Fig. 2b), consistent with the emerging post-transcriptional roles for eRNAs. The SF3B mutant exhibited altered junction usage in the cytoplasmic fraction under vehicle conditions, with 9 junctions differing from WT (FDR < 0.05), most showing increased usage in the mutant (7 of 9, Fig. 2c). Selinexor treatment in WT cells also altered junction usage in the cytoplasmic fraction, with 5 junctions differing from vehicle, including 3 increased and 2 decreased relative to vehicle (Fig 2d). In the cytoplasmic fraction, Selinexor responses differed modestly between genotypes, with 3 junctions showing genotype-dependent responses, including 2 with stronger Selinexor effects in WT and 1 in the SF3B mutant. In the nuclear fraction under vehicle conditions, the SF3B mutant differed from WT at 2 junctions, both showing increased usage in the mutant. Finally, a single junction showed localisation-dependent genotype differences in Selinexor response in WT cells. Together, these results indicate that eRNA splice usage varies with genotype, localisation, and environmental context.

To assess whether the splice junction perturbations observed following SF3B1 disruption reflected direct engagement of reconstructed eRNA transcripts with spliceosome-associated factors, we examined available crosslinking datasets for SF3B1. Analysis of published K562 iCLIP data (GEO: GSE247658)^14^ detected no reproducible SF3B1 interactions with the reconstructed eRNA transcripts, and interrogation of ENCODE eCLIP datasets identified only two reproducible SF3B1-eRNA interactions. Since SF3B1 recognises intronic branchpoints transiently as part of U2 snRNP rather than forming stable RNA contacts, the limited recovery of direct binding sites is consistent with its biochemical mode of action and does not contradict the junction-level perturbation effects observed above.

We therefore extended the analysis across the ENCODE eCLIP compendium (n=290)^15^ to determine whether reconstructed eRNAs show broader evidence of interactions with RNA-processing factors. Across all these datasets, multiple RNA-binding proteins (RBPs) interact with reconstructed eRNA transcripts, with the most frequently detected RBPs including CSTF2T, ILF3, IGF2BP1, HNRNPA1, FUS, TIA1, SRSF1, SAFB2, PPIL4, and NKRF, each interacting with several independent eRNAs (Fig. 2e). Many of these RBPs are known components of, or functionally linked to, spliceosome activity, RNA maturation, or transcript stability pathways ^11^. Thus, the reproducible detection of these interactions across multiple reconstructed eRNA transcripts supports the structural validity of the models and indicates that they are likely defined processing features rather than assembly artefacts.

### Multi-exonic eRNA splice junction usage differs between tumour and normal tissue

Aberrant RNA processing is a pervasive feature of cancer, reflecting disruption of spliceosome regulation, RNA localisation, and nuclear export pathways. Having identified these processes as determinants of eRNA junction usage, we asked whether similar regulatory patterns are evident in head and neck squamous cell carcinoma by analysing tumours alongside matched normal tissue. To enable this comparison, we first assessed whether reconstructed eRNA junctions could be detected in clinical RNA-Seq data. As poly(A)-selected sequencing is the dominant modality in clinical RNA-Seq, we tested whether these models remain observable in poly(A)-selected datasets. In 30 paired head and neck squamous cell carcinoma samples, 105 reconstructed splice junctions were testable for differential usage, of which ten (9.5%) differed between tumour and matched normal tissue (FDR < 0.05; Fig. 3a), including seven increased and three decreased in tumour.

**Figure 3.**
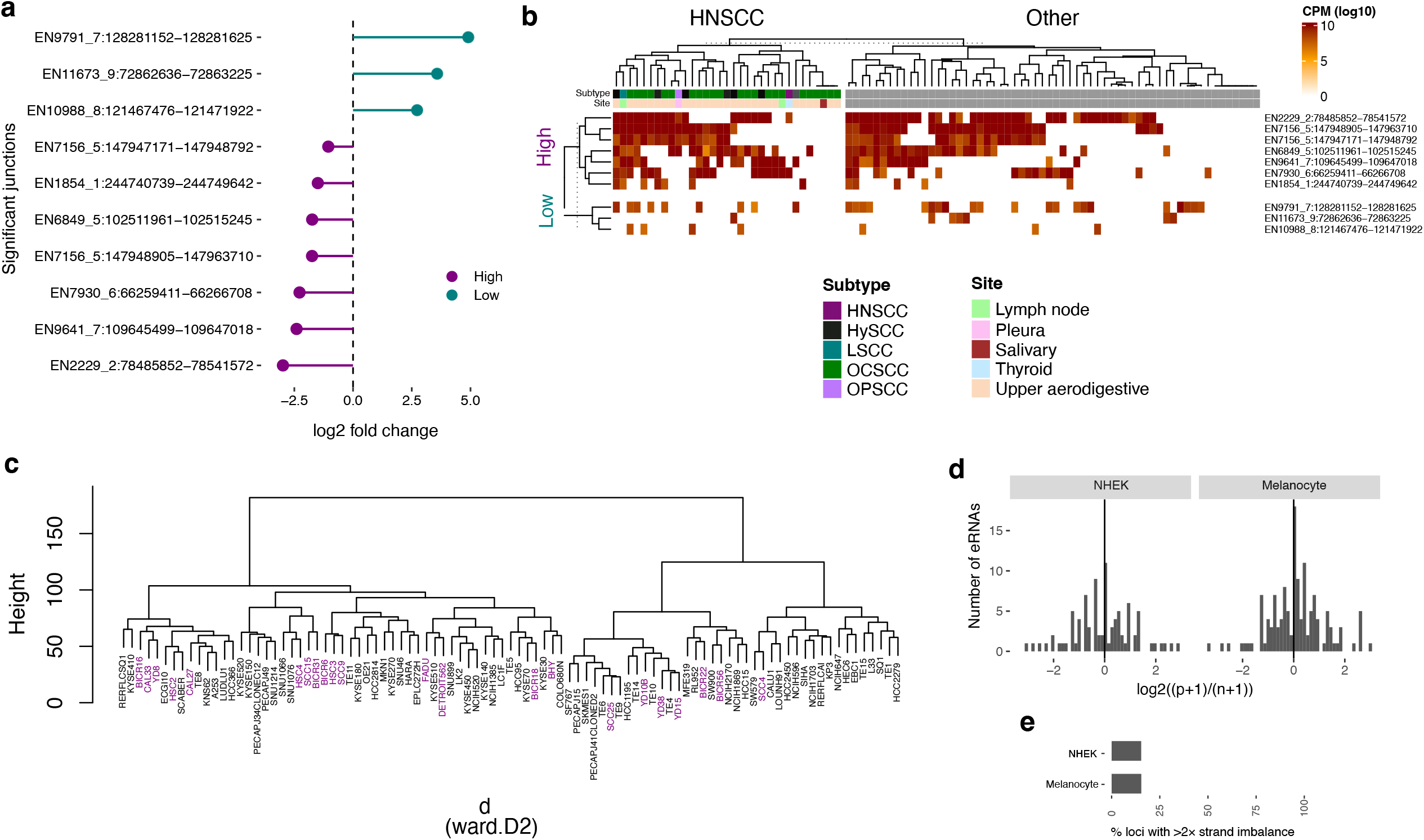
Tumour-associated enhancer RNA splice junctions and their representation in squamous carcinoma models. (a) Fold change of differential splice-junction usage between head and neck squamous cell carcinoma tumours and matched normal tissues. (b) Expression of differential eRNA splice-junctions across 93 squamous carcinoma cell lines from the CCLE dataset. Within the plot, SCC keys are OCSCC-oral cavity, OPSCC-Oropharynx, LSCC-Larynx, HySCC-Hypopharynx and HNSCC-head and neck SCC. (c) Hierarchical clustering of cell lines based on global eRNA splice-junction usage profiles, with squamous carcinoma cell lines marked in purple. (d) Distribution of strand asymmetry for reconstructed eRNAs in representative samples. Strand imbalance was calculated as log_2_((p+1)/(n+1)), where p and n denote counts assigned to the positive and negative strand components of each transcript, respectively. (e) Proportion of eRNAs exhibiting greater than twofold strand imbalance in each sample. Only loci with detectable signal were considered.

To assess whether these tumour-associated eRNA splicing events are observable in independent squamous systems that are amenable to mechanistic interrogation, we next examined their expression across a panel of squamous carcinoma cell lines from the Cancer Cell Line Encyclopaedia (CCLE; n = 94, poly(A)-selected). All ten junctions that were differentially used in head and neck tumours versus matched normal tissue were detectable within the CCLE squamous cohort (Fig. 3b), with junctions upregulated in tumours showing consistently high detection rate across the panel and those reduced in tumours remaining largely undetected. A small subset of cell lines derived from metastatic sites displayed modest deviations from this pattern, suggesting a potential role in disease progression that warrants further investigation. We therefore assessed global junction usage across the cohort. Hierarchical clustering revealed structured organisation of cell lines based on eRNA splicing profiles, including a compact subgroup enriched for head and neck carcinoma models (Fig. 3c). Together, these findings indicate that tumour-associated eRNA processing differences reflect lineage-related regulatory programmes and establish squamous carcinoma cell lines as tractable systems for investigating the regulation and functional consequences of tumour-associated eRNA processing changes.

### Analytical representation of bidirectional enhancers as paired RNA outputs

We next turn our attention to the broader class of eRNA transcripts, which comprise most of our reconstructed models (93% unstranded and unspliced) and are consistent with the established bidirectional and weakly processed nature of most eRNAs. Bidirectional transcription reflects the local recruitment of RNA polymerase II at enhancer regions without strong directional constraints. Consequently, transcription at these sites generate RNA outputs that diverge in opposite genomic directions. These transcripts are not reverse complements of a single molecule but represent distinct RNA species arising from opposite strands, each with its own sequence composition and structural potential. From an RNA perspective, a bidirectional enhancer therefore produces two separable transcriptional outputs rather than a single unified transcript. In practice, eRNA we collapse bidirectional transcription into a single measurement of enhancer output, reflecting the view of the enhancer as the primary regulatory unit rather than its individual eRNA products. While this representation captures overall enhancer engagement, it obscures the possibility that strand-specific RNAs may differ in persistence, localisation, or molecular interactions. Given that strand-of-origin alone generates non-identical RNA sequences capable of engaging distinct RNA-binding partners or decay pathways, such equivalence is unlikely to hold universally. For a non-coding RNA, function must arise from its RNA properties, making strand-specific differences in sequence or interactions especially informative.

To this end, we introduce a refinement to the unstranded and unspliced eRNAs in the catalogue by treating them as two linked strand-specific eRNA, each representing an independent RNA species while retaining a shared locus identity. We first addressed cases in which single exon transcripts overlapped regions already assigned to strand-resolved eRNA transcripts. Manual inspection showed that such transcripts most frequently trace intronic regions of stranded transcripts, and where they extend beyond annotated boundaries, read support is typically sparse. Because these overlaps would confound stranded read assignment, we excluded them from the catalogue and retained the strand-resolved annotations. For the remaining unstranded, unspliced transcripts, we generated strand-specific entries and append “p” or “n” to each identifier to denote positive or negative strand transcription. Strand-resolved quantification across an independent cohort of samples showed that paired enhancer outputs broadly track together yet are not uniformly symmetric at the RNA level (Fig. 3d). More than 12% of detected eRNAs exhibited greater than twofold strand asymmetry (Fig. 3e). While consistent with bidirectional transcription, this highlights the value of strand-resolved quantification and supports functional interrogation of eRNAs at the strand-specific molecular level.

We therefore provide the final filtered, strand-resolved GTF annotation (Supplementary Table S2) and junction BED file (Supplementary Table S3) as resources to support functional interrogation of eRNAs in the RNA space.

## Discussion

eRNAs have long been recognised as hallmarks of active enhancers ^1,4,16^, yet growing evidence indicates that many participate directly in gene regulation, both at the level of transcription and through emerging post-transcriptional mechanisms ^5,8,17^. Despite this shift in perspective, eRNAs remain primarily defined by genomic coordinates rather than by RNA molecules, limiting the ability to determine which transcripts mediate regulatory effects, how their sequence and structure recruit molecular partners, and how these interactions generate downstream outcomes. To address this limitation, we reconstruct enhancer transcription across diverse human tissues and cell types to establish a transcript-resolved framework for studying eRNAs as RNA molecules rather than genomic signals. Across 121 RNA-Seq datasets, we defined 19,484 transcripts spanning 18,740 loci and identified 1,538 splice junctions, nearly one-fifth of which were reproducibly detectable across independent perturbation and disease contexts. We show that some locus resolves into discrete RNA species with canonical splice architecture, conserved sequence features, and regulated usage patterns linked to localisation, mutation, and cellular perturbation, supporting their function within regulatory networks.

In independent validation datasets, including spliceosome perturbation in HeLa cells, perturbation in K562 cells, and head and neck tumour samples, reconstructed eRNA splice junctions showed broad detectability. In total, 400 of 1,538 junctions (26%) were observed in at least one validation dataset, indicating that a measurable fraction of the reconstructed catalogue is detectable beyond the samples used for assembly, despite the known context specificity of enhancer activity ^1^. A subset of these junctions also showed regulated usage across biological contexts. Across all contrasts tested in these validation datasets, 40 junctions (2.6% of the total catalogue; 14.6% of detectable junctions) exhibited significant differential usage, including responses to spliceosome perturbation, subcellular localisation, mutation status, drug treatment, or tumour state. These results indicate that splice junctions are reproducible features of some eRNA processing that can be detected and regulated across independent systems.

Several key observations emerge from this reconstruction that merit further consideration. First, enhancer loci overwhelmingly produced a dominant short transcript in our steady-state assembly, consistent with studies showing that enhancer transcription initiates focally but generates multiple short RNAs ^1,4,16^, of which only a subset persist long enough to be recovered in conventional RNA-Seq datasets. However, 1,379 transcripts contained multiple exons, and these displayed canonical splice-site motifs and reproducible junction usage (Fig 1e-g), indicating that at least a subset of eRNAs engage canonical RNA processing pathways characteristic of regulated noncoding RNAs. This contrasts with the prevailing assumption that eRNAs are predominantly unstructured or stochastic transcriptional outputs.

Second, splice-junction usage proved sensitive to biological context. Junction detection segregated by cell type and subcellular localisation, and perturbation experiments showed that junction usage responded to spliceosome moderation, nuclear–cytoplasmic partitioning, and inhibition of RNA export (Fig. 2a-d). These observations were notable because they demonstrate that eRNAs participate in the same layers of RNA regulation that govern other noncoding RNAs. In particular, the strong influence of localisation on junction usage suggests that eRNA processing is not solely dictated at the site of transcription but may continue to be shaped after transcription, confirming that some eRNAs participate in downstream RNA regulatory networks rather than remaining confined to nucleus.

A further unexpected observation was that reconstructed splice junctions remained detectable across diverse experimental platforms, including low depth single end sequencing and poly(A)-selected clinical datasets (Fig. 3a-d). Although eRNAs are often described as unstable and non-polyadenylated ^1^, these results confirm that a measurable subset persists long enough to be captured by conventional RNA-Seq and can exhibit regulated usage in disease contexts ^7,8^. Indeed, several had nearby polyA site (Fig. 1f), indicating presence of diverse functional classes.

Our transcript reconstruction relies on steady-state RNA-Seq signals, which preferentially capture stable and processed RNA molecules. Because many eRNAs are rapidly degraded, assemblies derived from these libraries inevitably enrich for eRNAs that persists long enough to be detected. Pooling multiple datasets improves sensitivity but biases models toward consensus structures that accumulate across conditions, potentially obscuring highly transient or context-restricted enhancer transcripts. Conversely, approaches that assemble transcripts within individual samples preserve condition specificity but often lack sufficient depth to resolve low-abundance eRNAs, resulting in fragmented or incomplete models ^5^. Some studies have attempted to address these limitations by reconstructing transcripts in systems where eRNA degradation is experimentally reduced, for example through exosome depletion or RNA stability perturbations. While such strategies increase recovery of unstable eRNA, they also alter the normal RNA lifecycle, potentially revealing intermediates or decay products that would not normally accumulate, and therefore risk conflating transient transcriptional by-products with physiologically stable eRNA species.

Another key observation directly related to the above point is that splice-junction discovery during read counting identified additional junctions beyond those incorporated into the reconstructed transcript models. This indicates that additional processed transcript exists beyond those captured in the current catalogue, particularly in disease contexts where RNA processing is perturbed. However, these additional junctions were typically supported by sparse reads and showed inconsistent detection across samples (Fig. 1k), raising uncertainty about their stability, reproducibility, and functional relevance. It therefore remains unclear whether such signals represent rare but genuine RNA isoforms, or transient processing intermediates. Resolving this ambiguity will require approaches that improve detection sensitivity while preserving molecular specificity. Long-read sequencing offers one potential solution for clarifying exon connectivity and validating rare isoforms, but the low abundance of many eRNAs means that such transcripts may remain difficult to capture without targeted enrichment. Strategies that selectively enrich eRNAs prior to sequencing could improve recovery, yet these approaches themselves will depend on prior transcript definitions to guide capture design. In this context, the transcript models presented here provide a starting framework for iterative refinement, enabling future studies to distinguish stable eRNA species from low-confidence transcriptional signals.

Despite these caveats, several observations argue that many reconstructed transcripts represent genuine regulated RNA species. Splice junctions showed reproducible detection across samples (206 of 1,538), broader recovery in perturbation experiments (456 of 1,538), and sensitivity to spliceosome perturbation. Junction usage varied with subcellular localisation (15 of 180) and tumour state (10 of 105), indicating that eRNA processing is responsive to canonical RNA regulatory pathways. These results do not imply that all enhancer transcripts are stable or functional, but they demonstrate that a subset behaves in ways consistent with regulated and regulatory noncoding RNA molecules.

Taken together, these findings show that the transcript-resolved models generated here provide a practical framework for studying eRNAs as defined molecular entities. By enabling transcript-level quantification and comparative analysis across contexts, this resource makes it possible to connect enhancer transcription to RNA processing, localisation, and regulatory interactions. As evidence continues to accumulate for roles of eRNAs in both transcriptional and post-transcriptional regulation, this work establishes a scalable platform for linking those functions to physiological and disease systems.

## Method

### Compilation of total RNA-Seq datasets

To reconstruct eRNA transcript models transcriptome-wide, we manually curated RNA-Seq datasets representing diverse human organs, cell types, and subcellular RNA fractions. We focused on released RNA-Seq datasets generated by the ENCODE consortium because their high sequencing depth, consistent library preparation, and uniform processing provide a robust foundation for pooled transcript assembly. In total, 121 RNA-Seq libraries representing 90 unique samples were compiled, including 54 cell line, 20 primary cell, 10 in-vitro differentiated cell, and 37 tissue samples. Majority of libraries were total RNA with rRNA depletion (n = 99), with smaller numbers of poly(A)-depleted (n = 8), poly(A)-selected (n = 8), and related fraction total RNA preparations (n = 6). A subset of libraries represented subcellular RNA fractions (n = 21), including cytosolic (n = 7), nuclear (n = 8), chromatin-associated (n = 2), nucleolar (n = 2), and nucleoplasm (n = 2) RNA preparations across K562 and HepG2 cells, enabling enhancer transcription to be captured across localisation states.

Detailed sample-level information is provided in Supplementary Table S2. BAM files (GRCh38/hg38) were downloaded directly from the ENCODE portal and used for transcript reconstruction.

### Construction of a human eRNA pan-transcriptome

To reconstruct eRNA transcript models, we performed de novo transcript assembly of RNA-Seq alignments restricted to curated enhancer loci. Reads were filtered to retain only those overlapping a curated set of eRNA coordinates (48,766 loci), thereby reducing signal from gene bodies and promoters while preserving transcript structure within enhancer regions. To maximise sensitivity for low-abundance eRNAs, filtered BAM files from all samples (n = 123) were pooled into a single alignment using samtools (v1.19.2) ^18^. De novo transcript assembly was then performed on the merged BAM using StringTie (v3.0.3) ^19^ with a minimum transcript length threshold of 100 nt (-m 100). For pilot analyses, transcript assembly was also performed independently on full per-sample BAM files using identical parameters. To minimise contamination from gene-associated transcription, reconstructed transcripts overlapping RefSeq gene bodies were removed, and models exceeding 100 kb were excluded. The final transcript set (Supplemental Table S2) was assigned locus-level identifiers (EN) and cross-indexed to the originating enhancer coordinates through the GTF attribute eRNAkit.

### Poly(A) site proximity analysis

We first defined the 3′ end of each transcript as the end position of the terminal exon in the direction of transcription, and the 5′ end as the start position of the first exon. End positions shared by multiple transcripts were collapsed to unique intervals (1 nt). We extracted poly(A) sites from PolyA Atlas (GRCh38) that within 500 nt in the correct orientation using BEDtools closest ^20^. For 3′ ends, we restricted candidate poly(A) sites to downstream positions on the same strand (parameters: -s -iu -D a), and for 5′ ends we restricted candidates to upstream positions on the same strand (-s -id -D a). To estimate local background poly(A) site density, we repeated the same directional queries on the opposite strand (-S), generating orientation-mismatched controls. We then used the resulting count distributions from strand-matched and strand-reversed comparisons to assess enrichment of reconstructed 3′ termini near annotated cleavage and polyadenylation sites.

### Splice-site motif analysis

For each junction, we defined the donor site at the exon–intron boundary of the upstream exon (the last exonic base on the + strand or the first exonic base on the − strand) and the acceptor site at the intron–exon boundary of the downstream exon (the first exonic base on the + strand or the last exonic base on the − strand). We then extracted strand-aware sequences corresponding to established splice-site motif models: donor windows spanning 3 nt of exon and 6 nt of intron (−3 to +6 relative to the donor junction) and acceptor windows spanning 20 nt of intron and 3 nt of exon (−20 to +3 relative to the acceptor junction) ^21^. Identical donor or acceptor windows arising from multiple transcripts were collapsed into unique sequences for downstream motif analysis using MEME (v5.5.9) online with default parameters in classic discovery mode.

### Validation splice junctions using independent RNA-Seq datasets

To validate reconstructed eRNA splice junctions in independent datasets, we analysed publicly available RNA-Seq studies not associated with the assembly cohort (GEO: GSE255179, GSE72055, GSE285603) ^12,13,22^, together with a panel of 93 squamous carcinoma cell lines from the Cancer Cell Line Encyclopaedia (CCLE) dataset ^23^. Detailed sample-level information is provided in Supplementary Table S3. Raw RNA-Seq reads corresponding to these datasets were obtained from SRA and processed using our established pipeline ^24^. Reads were quality filtered (Q < 20) and adapter trimmed using Trimmomatic (v0.39) ^25^, then aligned to the human reference genome (hg38) using HISAT2 (v2.1.0) with default settings ^26^. Reads were filtered to retain only those overlapping the reconstructed transcript coordinates to enable effective comparison between reconstructed junctions (KNOWN) and background splice signals (NOVEL). This filtering step is not required for routine use of the GTF annotation and was applied solely for benchmarking purposes. Counts of reads overlapping junctions were quantified using featureCounts (Subread v2.1.1) ^27^ against reconstructed eRNA exon-exon junctions (.BED) created from the transcript models (Supplemental Table S2). The usage levels were normalised as counts per million (CPM). We recommend performing this normalisation exclusively on eRNA-derived counts, preferably restricted to KNOWN junctions, to minimise influence from abundant mRNA splice junctions.

Differential splice-junction usage between experimental groups was assessed using the Limma framework (v3.64.3) ^28^. Log^2^-transformed CPM values were used directly for modelling. Linear models were fitted for each junction, and empirical Bayes moderation was applied to obtain moderated statistics. P values were adjusted for multiple testing using the Benjamini–Hochberg method, with FDR < 0.05 considered significant.

### Validation of unstranded transcript strand splitting

We analysed publicly available RNA-Seq studies not associated with the assembly cohort or used in the splice junction validation (ENA: E-MTAB-5403) ^29^. Preprocessing and alignment were performed as described above for the splice junction validation set. eRNA transcript counts were quantified using HTSeq (v0.11.1)^69^ against reconstructed eRNA transcripts with parameters -s yes and -m intersection-strict.

### RNA binding protein iCLIP and eCLIP overlap analysis

To assess eRNA–RBP interactions with reconstructed eRNA transcripts, we analysed publicly available crosslinking datasets. We downloaded pre-processed merged peak files from SF3B1 iCLIP in K562 cells (GEO: GSE247658) and from the ENCODE eCLIP compendium and used them directly. We identified overlaps between crosslinking peaks and reconstructed eRNA transcript loci using the R GenomicRanges package ^30^. We summarised detected overlaps per protein to quantify recurrent associations between RNA-binding proteins and reconstructed eRNA transcripts.

### Statistical analysis

Unless otherwise stated, differences between two groups were assessed using the Wilcoxon rank-sum test, whereas comparisons involving more than two groups were evaluated using the Kruskal–Wallis test, as specified in the figure legends. Statistical significance was defined as p < 0.05 for single comparisons and as FDR < 0.05 for multiple testing.

## Data availability

All sequencing datasets analysed in this study are publicly available through the original repositories from which they were obtained (ENCODE, GEO and SRA; accession numbers provided in Supplementary Tables S1). The reconstructed eRNA transcript annotation generated in this work is provided as a GTF file in the eRNAkit repository, as data slot in eRNAKitDB.rds (https://github.com/AneneLab/eRNAkit), and included as Supplementary Data S2. Associated junction annotation BED is available through the same repository and database entry.

## Author contributions

C.A.A. conceived the project, curated the datasets, and performed the bioinformatic analyses. N.B., R.K., E.I., J.B., and C.A.A. analysed and interpreted the data. C.A.A. supervised the project and wrote the manuscript. All authors edited and approved the final manuscript.

